# Differential enrichment of retinal ganglion cells underlies proposed core neurodegenerative transcription programs

**DOI:** 10.1101/2024.10.21.618927

**Authors:** Trent A. Watkins

## Abstract

In a published Correction^1^, a revised analysis updated two “core transcription programs” proposed to underlie axon injury-induced retinal ganglion cell (RGC) neurodegeneration. Though extensive, the Correction purported to leave the two principal conclusions of its parent study^2^ unaltered. The first of those findings was that a core program mediated by the Activating Transcription Factor-4 (ATF4) and its likely heterodimeric partner does not include numerous canonical ATF4 target genes stimulated by RGC axon injury. The second was that the Activating Transcription Factor-3 (ATF3) and C/EBP Homologous Protein (CHOP) function with unprecedented coordination in a parallel program regulating innate immunity pathways. Here those unexpected findings are revealed to instead reflect insufficient knockout coupled with differences in RGC enrichment across conditions. This analysis expands on the published Correction’s redefinition of the purported transcription programs to raise foundational questions about the proposed functions and relationships of these transcription factors in neurodegeneration.

## Introduction

Transcriptional programs activated in response to axon injury are important regulators of neuronal survival and axon regenerative potential. Upregulation of Activating Transcription Factor-3 (ATF3) following axonal injury signaling, for example, contributes to a broad range of transcriptional changes within adult sensory neurons after peripheral nerve injury and is necessary for efficient axon regeneration^3^. ATF3 functions, at least in part, in coordination with another stress-activated basic leucine-zipper (bZIP) transcription factor (TF), c-Jun, a heterodimeric partner of ATF3 that is also required for axon regeneration^4^. Neuronal ATF3 and c-Jun are also activated, along with many other TFs, by some forms of CNS axon injury, such as optic nerve crush, though regeneration of the damaged retinal ganglion cell (RGC) axons is thwarted by intrinsic and extrinsic barriers. The transcriptional injury response in that context results in widespread neuronal apoptosis.

Several studies have attempted to identify the TFs that contribute to RGC neurodegeneration after axon injury. Using conditional knockout (cKO) mice, c-Jun has been identified as a primary neuron-autonomous promoter of RGC apoptosis^5^. Independent *in vitro* siRNA screening not only confirmed a central role for c-Jun but also found contributions from ATF2, Sox11, and MEF2A^6^. Also implicated in RGC death are the Integrated Stress Response (ISR) effectors Activating Transcription Factor-4 (ATF4) and C/EBP Homologous Protein (CHOP)^7,8^. Conditional knockout of ATF4 or its upstream activator PKR-like Endoplasmic Reticulum Kinase (PERK) abrogates the potent injury-induced upregulation of numerous known ATF4 target genes and confers significant but incomplete neuroprotection^8^. Germline knockout of *Ddit3*, the gene encoding CHOP, is mildly neuroprotective after optic nerve crush but has minimal impact on the retinal transcriptional injury response^5,9^. Highlighting the complex interrelationships amongst these and other potential bZIP heterodimeric partnerships, injury responses include transcriptional changes that are: (1) dependent on both c-Jun and ATF4; (2) dependent on one but not the other; and (3) dependent on neither^8,9^. ATF4, like c-Jun, also contributes to RGC axon regenerative potential^8^.

As one of three related studies featured in an accompanying Commentary^10^, Tian *et al*. reported *in vivo* CRISPR screening of 1,893 transcription factors for their contributions to RGC neurodegeneration and axon regeneration after optic nerve crush^2^. gRNA pools targeting ten individual TFs were found to confer neuroprotection, including gRNAs intended to knock out ATF3, CHOP, ATF4, and a known heterodimeric partner of ATF4, C/EBPγ. Perhaps surprisingly, c-Jun was not among the reported “hits” of the screen. This screen was complemented by a chromatin accessibility study (ATAC-seq) in which DNA “footprints” were scanned for TF binding motifs to infer which TFs might be bound to expressed genes. A third leg of the study used RNA-seq of FACS-enriched, gRNA-expressing RGCs at 3 days post-crush (3dpc) to identify direct and indirect transcriptional consequences of ATF3, ATF4, C/EBPγ, and CHOP knockout. The convergent results of these approaches were proposed to map out two parallel, partially overlapping, neurodegenerative transcription programs, one mediated by ATF3/CHOP and the other by ATF4/C/EBPγ^2^. Surprisingly, the proposed programs included exceptionally few of the major expression changes well-documented to be injury-induced in the same study and many others^2,5,8,11,12^. Moreover, they were quite distinct from the known transcription programs mediated by these TFs across a variety of cellular stress contexts^3,13–16^. Among many other differences, the pronounced overlap between ATF3 and CHOP function, in parallel to ATF4, represents a departure from the canonical ATF4-ATF3-CHOP cascade^17^. Together, the findings suggested novel relationships among these TFs and their target genes unique to this model of CNS axon injury.

A second published Correction to Tian *et al*. (2022) updated multiple figures, supplemental figures and tables, as well as the underlying analyses of the RNA-seq data posted to the Gene Expression Omnibus (GEO) as GSE190667^1^. This *Matters Arising* evaluates the impacts of that Correction and assesses the extent to which the amended data and analyses support the central conclusions of the manuscript. This evaluation reveals that the Correction substantially alters the criteria and constituents of the “core transcription programs,” revealing underlying deficiencies in efficacy of knockout of the targeted TFs and in the control of batch effects. These limitations resulted in assignment of numerous non-RGC transcripts to the proposed RGC-autonomous “transcription programs” and render the findings of neuroprotection upon gRNA expression uninformative for clarifying the functional roles and relationships of these TFs in neurodegeneration.

## Results

### Shifted inclusion criteria weaken the case for programs coordinated by pairs of TFs

The centerpiece of a manuscript entitled “Core transcription programs controlling injury-induced neurodegeneration of retinal ganglion cells,” is, one may infer, the proposed “programs,” which are depicted in Tian *et al*.’s Figure 5 and Table S6. Table S6 lists all genes included in each of three proposed programs: ATF3/CHOP, ATF4/C/EBPγ, and “Common” (*i*.*e*., dependent on all four of these TFs). Tian *et al*.’s Figure 5A diagrammatically represented a subset of those genes proposed to participate in the top biological pathways uncovered in the analysis, such as “TLR Signaling and Processing” for the ATF3/CHOP program and “Autophagy” for the ATF4/C/EBPγ program. Tian *et al*.’s Figure 5B shows the footprint signals for two selected genes from each of the three programs, one serving as an exemplar of a gene bound and positively regulated by that program’s TFs and the other serving as an exemplar of a gene bound and negatively regulated by that program’s TFs. A Correction revised the programs proposed in the original publication, including three of the six exemplar genes of Tian *et al*.’s Figure 5B. The nature and significance of those updates can be determined by comparing the overlap between the original and revised analyses^1,2^. Quantification reveals that 20-50% of the genes in each of the original three Table S6 “programs” have been excluded in the Correction, along with 42% of the genes in Tian *et al*.’s original Figure 5A (**Figure 1A-D**). Unexpectedly, the revised Table S6 programs nevertheless expanded by 153%-379% on account of newly included genes, with ten of these new genes added to Tian *et al*.’s central Figure 5A. These results imply that the Correction may have involved a significant redefinition of the proposed model of parallel neurodegenerative transcription programs mediated by these pairs of TFs.

**Figure 1.**
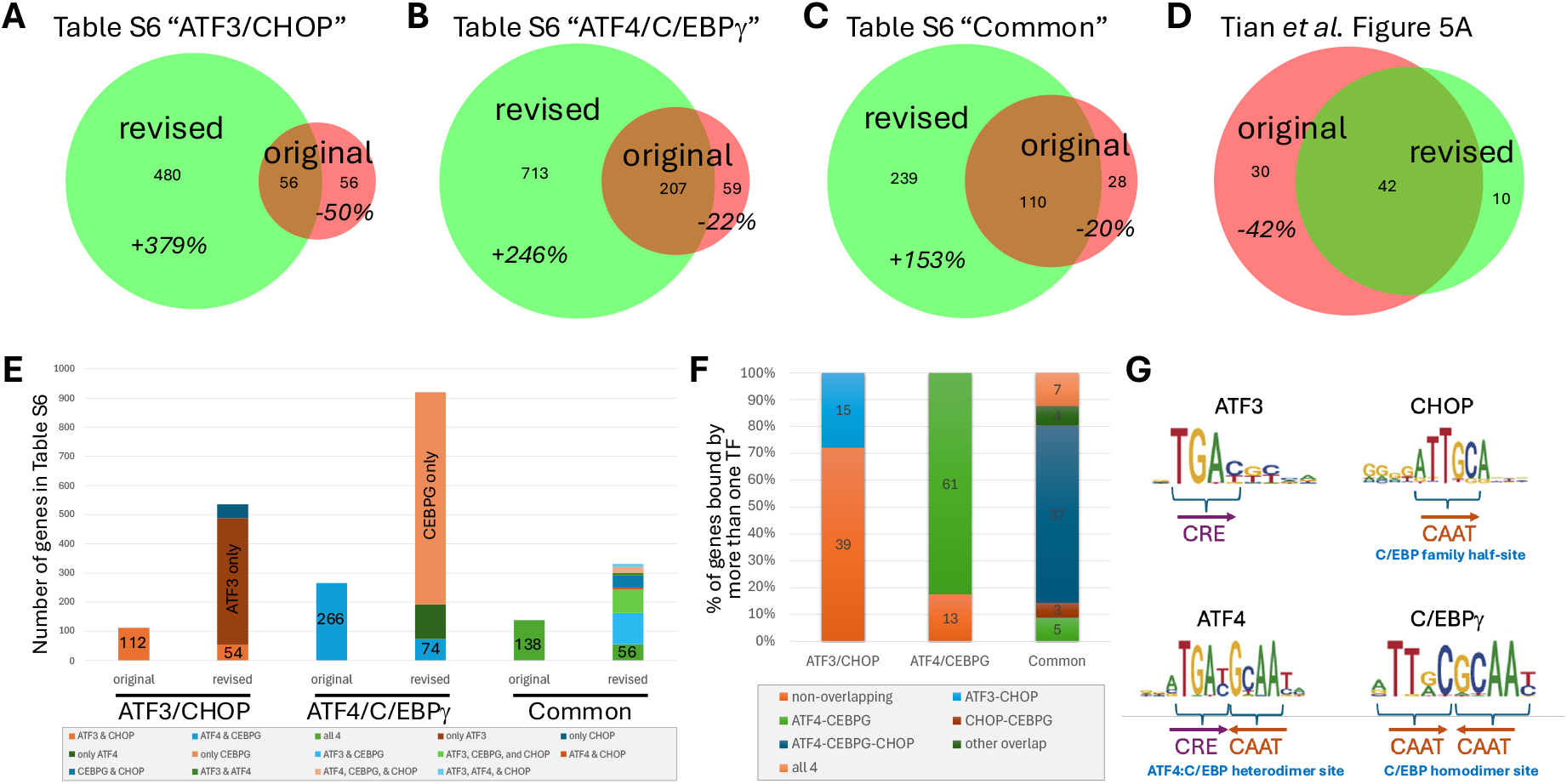
Shifted inclusion criteria weaken the case for “programs” coordinated by pairs of TFs. (A-D) Venn diagrams of genes included in Tian *et al*.’s original and revised Table S6 and Figure 5A proposed “transcription programs” reveal the extent of genes lost in the Correction, as well as numerous genes added. (E) Counts of genes in the original and the revised Table S6 exhibiting “binding” by multiple TFs inferred by DNA footprinting, with co-“bound” genes serving as a defining criterion in the originally proposed programs but representing a small fraction of the revised programs. (F) Quantification of overlap of inferred binding sites in the revised Table S6 programs amongst the remaining genes reported to have more than one TF site. (G) Breakdown of motifs used in DNA footprinting analysis shows that the motif used for ATF4 is an amalgam of the CRE half-site preferred by ATF4 and the CAAT half-site preferred by C/EBP family members. All data has been quantified from Tian *et al*.’s original and revised Figure 5A and Table S6, which reported the constituent genes of the proposed core neurodegenerative transcription programs.

It is important to understand any differences in inclusion criteria that might account for the simultaneous compression of Figure 5A and expansion of Table S6 proposed “core transcription programs.” A seeming premise of the Tian *et al*. manuscript was that the TF pairs that seemed to show common functionality in neurodegeneration would also exhibit co-regulation of common transcripts. Amongst those co-regulated transcripts, genes for which DNA footprinting suggested possible binding by both TFs were proffered as their direct “transcription program”^2^. That premise suggested that inclusion criteria for the programs of Table S6 might include: (1) inferences from DNA footprinting that a gene is bound by all the relevant TFs that define each program; and (2) significant effects of all gRNAs targeting the relevant TFs. Evaluation of the original and revised Tables S6 shows that the first of these criteria was essential for inclusion in the original Table S6 but now represents just a small fraction of the genes included in the revised Table S6. The number of genes in the revised “programs” meeting this previously essential criterion is now reduced by 51% for ATF3/CHOP, by 72% for ATF4/C/EBPγ, and by 61% for the “Common” program (**Figure 1E**). The programs delineated in the revised Table S6 nevertheless expanded with numerous genes for which DNA footprinting-inferred “binding” of a single TF was sufficient for inclusion, with 89.9% of genes in the revised ATF3/CHOP program now listed in Table S6 as bound only by ATF3 or only by CHOP and 92.0% of genes in the ATF4/C/EBPγ program now listed as bound only by ATF4 or only by CEBPG. Notably, 100% of genes in the revised Table S6 meet the second criterion of exhibiting significant effects of gRNA at FDR<0.1 for each listed TF binding site, a departure from the original analysis for which *p*<0.05 was in most cases sufficient for inclusion in the program, including transcripts reported at FDR>0.2 for one or more of the relevant TFs (**Figure 1E**).

This evaluation reveals a significant shift in the defining features of the proposed “core programs” between the original manuscript and the latest Correction. Whereas the originally published analysis placed emphasis on genes co-”bound” by the relevant TFs (with minimal criteria for gRNA effects), the revised analysis appears to do the opposite. It largely eschews the common binding and co-regulation criteria of the original model in favor of any transcript that exhibits “binding” by one of the TFs (inferred by DNA footprinting), along with exhibiting significant effects (at FDR<0.1) upon expression of its corresponding gRNA. This raises important questions about the degree to which the underlying data are consistent with the proposed coordinated regulation by the TF pairs presented in the model.

### To what extent do TFs co-regulate the transcripts within their shared programs?

One mechanism that can dictate the degree of co-regulation by pairs of TFs is whether they function as a heterodimeric pair. Evaluating overlap between inferred TF binding loci in Tian *et al*.’s DNA footprinting analysis can therefore inform assessment of the hypothesized coordinated TF action^1^. Among the revised Table S6 “program” genes that have inferred binding sites for more than one TF, fewer than 30% of the ATF3/CHOP program genes are reported to exhibit overlapping binding sites for ATF3 and CHOP, inconsistent with a prominent role of ATF3-CHOP heterodimers in their proposed regulation of common target genes (**Figure 1F**). In contrast, greater than 80% of the co-”bound” ATF4/C/EBPγ program genes have overlapping inferred binding sites consistent with ATF4-C/EBPγ heterodimerization. Finally, approximately 2/3 of “Common” genes include at least one inferred binding site for ATF4-CHOP-C/EBPγ plus one or more separate sites for ATF3. These bioinformatic inferences by themselves provide minimal insight, as they are largely predetermined by features of the motifs used to search for potential TF bindings sites. For instance, the motif that was used to search for “ATF4” binding sites is actually a union of the half-site preferred by ATF4 and the half-site preferred by C/EBP family members, including C/EBPγ and CHOP (**Figure 1G**)^18^. It is therefore unsurprising that inferred “ATF4” sites would frequently overlap with inferred C/EBPγ sites. This result nevertheless highlights an important aspect of the underlying biology. Because ATF4 and C/EBPγ typically function together as heterodimeric partners in a variety of contexts, and because 92.9% of RGC-expressed genes known to be co-bound and co-regulated in another model of cellular stress^19^ are also upregulated following optic nerve crush, it is expected that many target genes, such as *Sesn2, Pck2, Herpud1*, and *Slc7a11*, should, in a dependable data set, exhibit similar dependency on these two TFs. In contrast, because ATF3 and CHOP instead target distinct regulatory sites, the effects of knockout of each are expected to be more divergent.

To evaluate whether the published data of Tian *et al*. are consistent with these basic expectations, pair-wise plots were generated of the reported effects of each TF gRNA on the genes within each proposed “program” delineated in revised Tian *et al*.’s Table S6 and Figure 5A^1^. In stark contrast to the DNA footprinting prediction that ATF3 and CHOP largely target distinct regulatory loci, the effects of ATF3 gRNA and CHOP gRNA on Table S6 program genes were by far the most concordant with one another of any TF pairs across all three programs, with strikingly high R^2^ coefficients of 0.86-0.97 (**Figure 2A-C**). Also contrary to expectations, the effects of gRNA targeting ATF4 and gRNA targeting its likely heterodimeric partner CEBPG on Table S6 genes were the most divergent from one another, exhibiting little consistent commonality, especially within the proposed ATF4/C/EBPγ program (**Figure 2B**). This discordance extends to genes for which the inferred ATF4 and C/EBPγ binding sites are overlapping according to Tian *et al*.’s Table S6, such as *Akna, Car2, Clcn7*, and *Fezf2*, each with reported effects of ATF4 gRNA in the opposite direction of effects of CEBPG gRNA. These results introduce an unexpected paradox. How might ATF3 and CHOP gRNA exhibit extraordinarily similar effects on the proposed direct targets of those TFs, despite expected binding to distinct regulatory sites, whereas ATF4 gRNA and CEBPG gRNA effects share the least in common, despite a predominance of overlapping inferred binding sites suggestive of functional heterodimerization?

**Figure 2.**
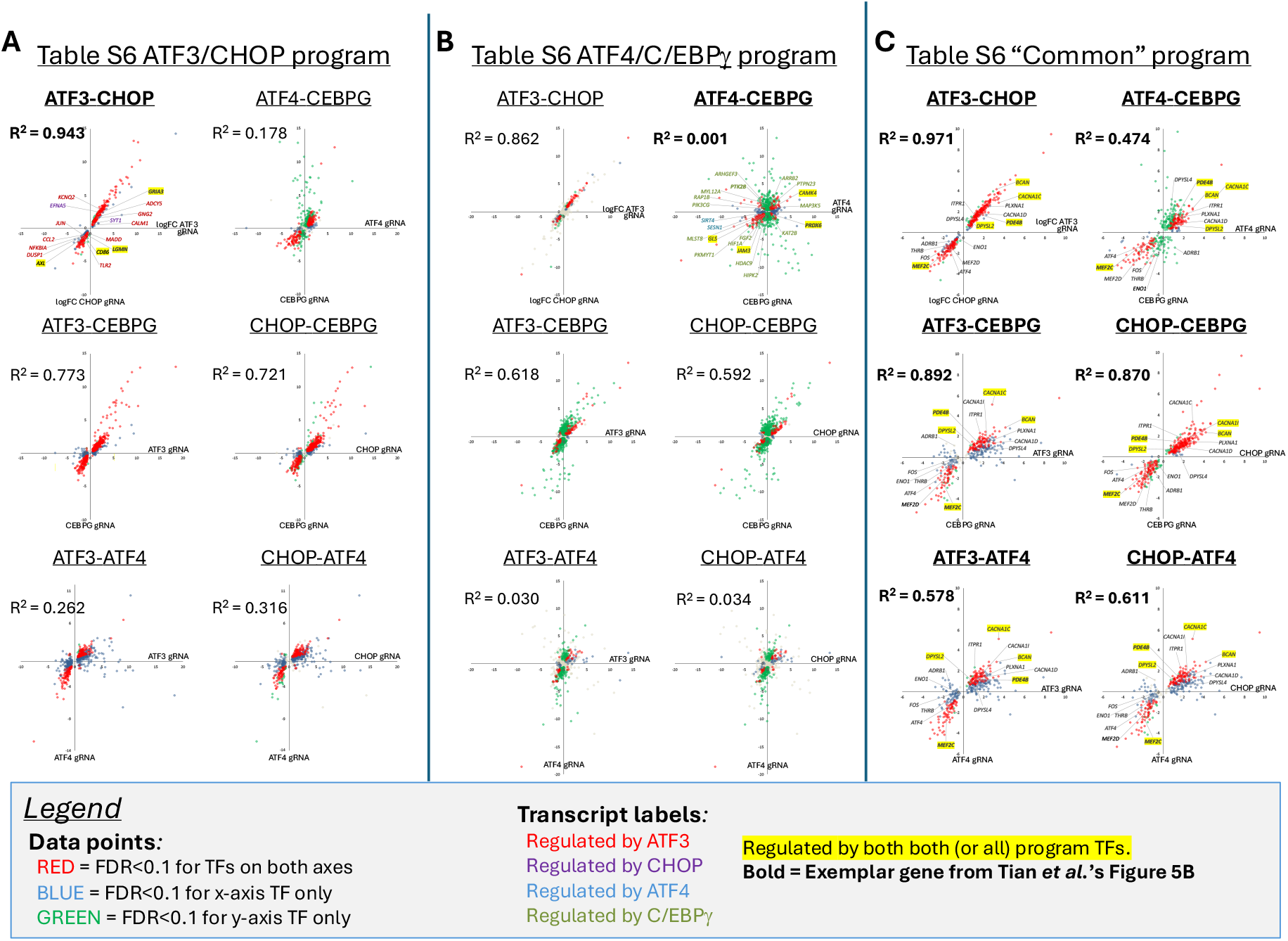
Unexpected extent of co-regulation within core transcription programs. For the genes in each of the three proposed programs reported in Tian *et al*.’s revised Table S6 – (A) ATF3/CHOP, (B) ATF4/C/EBPγ, and (C) Common to all 4 TFs – pair-wise plots were generated of the differential expression values reported in Tian *et al*.’s revised Table S5 and GSE190667_rna.DEX. Each graph plots the effects of one gRNA (log_2_ fold-change, or logFC) versus the effects of another gRNA (logFC) on the proposed program genes, with data points color-coded to represent which logFC values were reported as significant at FDR<0.1 across conditions. For the most relevant pair-wise plot(s) in each program (*e*.*g*., ATF3 vs. CHOP effects for the ATF3/CHOP program), data points are labelled with transcript names for genes that are also presented in Tian *et al*.’s Correction Figure 5, color-coded to represent the connections depicted in that published diagram. (A total of 13 genes are highlighted to indicate connections to all the relevant TFs within each program, down from 46 that met this criterion in the original Figure 5A.) Strangely, despite distinct inferred binding sites on most proposed program genes, the effects of ATF3 gRNA and CHOP gRNA were most similar to one another across all three programs, whereas ATF4 gRNA and CEBPG gRNA effects were most divergent despite their likely function as a heterodimer.

### Weak relationships between injury-responsiveness and gRNA effects

Further puzzling inconsistencies emerge from evaluation of the injury-responsiveness of Tian *et al*.’s proposed program genes. Independent studies of optic nerve injury have found, unsurprisingly, that the effects of conditional knockout of c-Jun or ATF4 are overwhelmingly to diminish injury-induced expression changes^8,9^, consistent with these injury-activated TFs acting as the mediators of those transcriptional stress responses. Plotting the reported gRNA effects of Table S6 genes against the 3dpc injury time course data of Tian *et al*.’s GSE184547 therefore provides an opportunity to assess how Tian *et al*.’s proposed core programs relate to the transcriptional injury response thought to underlie neurodegeneration^1,2^. Within the ATF3/CHOP program, many transcripts exhibit the expected inverse relationship between injury-induced expression changes and effects of ATF3 gRNA and/or CHOP gRNA (**Figure 3A**). For example, *Gria3*, plotted in Quadrant II, was reported to be significantly downregulated by injury (negative log_2_ fold-change, or logFC, at 3dpc compared to uninjured) but this downregulation was suppressed by ATF3 gRNA (positive logFC comparing ATF3 gRNA to non-targeting control gRNA, both at 3dpc). Despite these trends, other proposed program genes do not appear to be injury-responsive, and some show discordant effects. Three of the six genes presented in Tian *et al*.’s Figure 5A in support of “Neuroinflammation” as the top biological pathway of the ATF3/CHOP program - *Axl, Cd86*, and *Ccl2* - are found in Quadrant III, reported to be downregulated by 3dpc after injury (negative logFC) yet even more strongly downregulated after injury in the ATF3 gRNA condition (negative logFC). This observation, which is corroborated by Tian *et al*.’s own Table S3A report of these genes as “Down” in both expression and chromatin accessibility after injury, is incongruent with injury-stimulated ATF3 mediating that aspect of the response. This incongruence is even more pronounced within the proposed ATF4/C/EBPγ program, with no clear relationship between injury responsiveness and gRNA effects (**Figure 3B**). The transcript *Gls*, for example, is downregulated by injury, though it is also reported to have a common ATF4/C/EBPγ binding site and to be positively regulated by both of those injury-activated TFs. Additional incongruent results include *Mef2d* of the “Common” program, reported as downregulated following injury despite proposed positive regulation by injury-activated ATF3 and C/EBPγ (**Figure 3B**). These findings are not consistent with the proposed TFs mediating these injury-responsive transcriptional changes and raise the possibility that the reported differentially expressed genes (DEGs) reflect something other than knockout of the targeted TFs.

**Figure 3.**
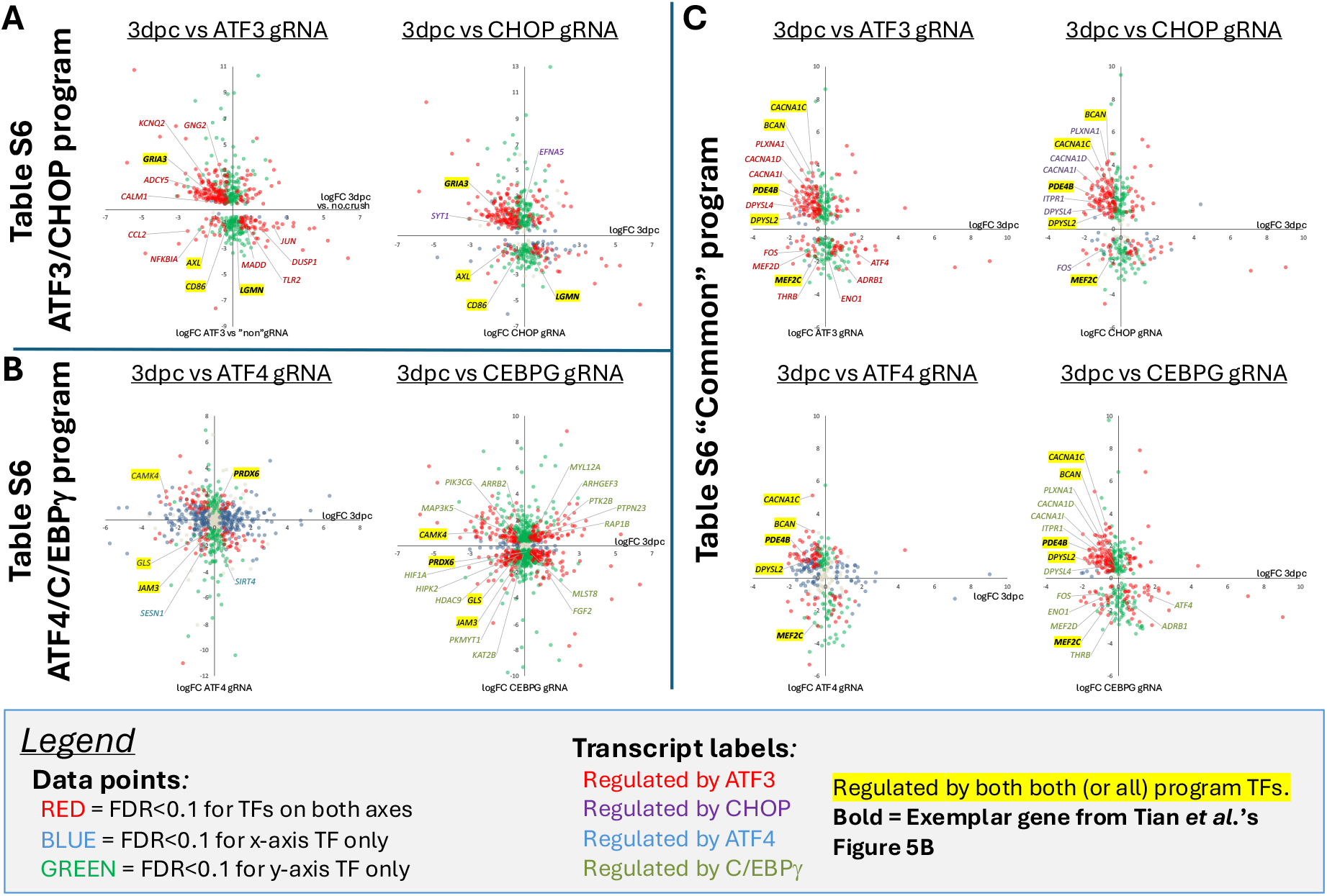
Few proposed program genes are injury-responsive coherent with gRNA effects. For the genes in each of the three proposed programs reported in Tian *et al*.’s revised Table S6 – (A) ATF3/CHOP, (B) ATF4/C/EBPγ, and (C) Common to all 4 TFs – pair-wise plots were generated of the differential expression values reported in Tian *et al*.’s revised Table S5 / GSE190667_rna.DEX and an independent DESeq2 analysis of Tian *et al*.’s GSE184547 RNA-seq data set reporting RGC expression changes at 3 days post-crush (3dpc) injury. Each graph plots the reported effects of a gRNA (logFC) versus the reported injury-induced expression change (logFC), with data points color-coded to represent which logFC values were significant at FDR<0.1. For genes also presented in Tian *et al*.’s Figure 5 as regulated by the relevant TF, data points are labelled with transcript names (with yellow highlight indicative of connections to all relevant TFs within a program). Numerous data points appear in Quadrants I and III, such that injury-induced changes are amplified, rather than mitigated, by gRNA expression, inconsistent with the interpretation that the targeted TF mediates that injury response.

### The gRNAs were largely ineffective against their intended TF targets

Though not an absolute prerequisite for CRISPR-mediated knockout, nonsense-mediated decay (NMD) of the targeted mRNA is highly correlated with effective disruption of function^20^. Probing Tian *et al*.’s RNA-seq data for evidence of NMD of the gRNA-targeted TFs can therefore inform the likelihood of functional knockout (**Figure 4A**). As expected from numerous studies, mRNA for each of the “core” TFs exhibited upregulation by RGCs following injury, with *Atf3* mRNA increasing by greater than 100-fold by 3dpc in Tian *et al*.’s time course data set reported in Table S2 and GEO: GSE184547^2^. In the Table S5 and GEO: GSE190667 data set reporting effects of gRNAs at 3dpc, *Atf3* mRNA exhibited 32% reduction in the CEBPG gRNA condition and 70-75% reduction upon expression of gRNAs targeting ATF3 or CHOP. Given the magnitude of *Atf3* upregulation upon injury, this moderate reduction might be expected to still leave *Atf3* expression at least 20-fold above its uninjured baseline. Though all four gRNA treatment conditions resulted in modest reduction of *Atf4* expression values compared to control, the 45% reduction of *Atf4* mRNA in the ATF4 gRNA was the least suppressed of the four conditions. RNA-seq reported *Cebpg* to be reduced by a modest 27-34% in the ATF3, ATF4, and CHOP gRNA conditions, but not significantly altered in the CEBPG gRNA condition. *Ddit3*, the mRNA encoding CHOP, was unaffected by CHOP or CEBPG gRNA, elevated by 42.6% by ATF4 gRNA, and decreased by 23% by ATF3 gRNA. Together these data suggest the possibility of moderate knockout *Atf3* in ATF3 gRNA and CHOP gRNA conditions but no discernable pattern of substantial on-target reduction of other targets.

**Figure 4.**
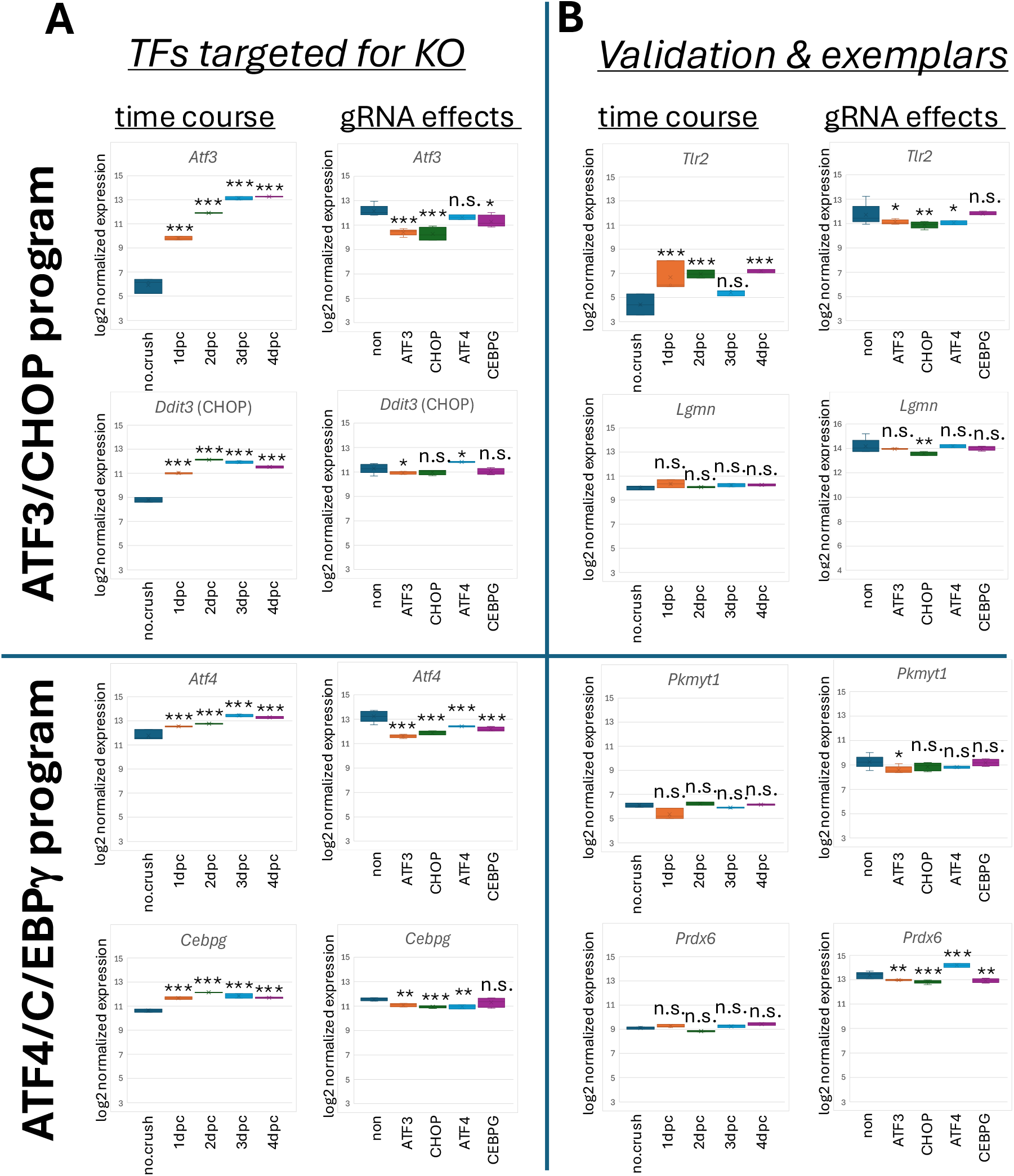
Limited evidence for substantial knockout of targeted TFs. (A) Box-and-whisker plots of injury-induced expression changes of the four proposed core TFs and the effects of gRNAs targeting these TFs. (B) Box-and-whisker plots of injury-induced expression changes of two of Tian *et al*.’s exemplar program target genes (*Lgmn, Prdx6*) and the two proposed validation genes (*Tlr2, Pkmyt1*) and the effects of gRNAs targeting the core TFs on these genes. All data are from independent OneStopRNA-seq DESeq2 analyses of Tian *et al*.’s GSE184547 (injury time course) and GSE190667 (gRNA effects) RNA-seq data sets. GSE184547: *n*=3 for “no.crush” control, 1dpc, and 3dpc time points; *n*=2 for 2dpc and 4dpc. GSE190667: *n*=5 for “non”-targeting control, ATF3 gRNA, and CHOP gRNA conditions; *n*=4 for CEBPG gRNA; *n*=2 for ATF4 gRNA. *p(adj)<0.1; **p(adj)<0.01; ***p(adj)<0.001; n.s.=not significant.

Ineffective knockout is consistent with the absence of known injury-induced ATF4-dependent target genes amongst the reported DEGs^8,13,19^. To determine if other data provided by Tian *et al*. might be consistent with functional knockout, plots of GSE184547 and GSE190667 data were also generated for: (1) two genes highlighted in Tian *et al*.’s Figure 5B as exemplars of the co-bound, co-regulated genes of the proposed programs, *Lgmn* and *Prdx6*; and (2) two genes purported in Tian *et al*.’s Figure S5 to be validated by immunohistochemistry (IHC) as injury-responsive and disrupted by corresponding gRNAs, *Tlr2* and *Pkmyt1* (**Figure 4B**). These purported exemplars and validation genes might be expected to provide the clearest support for functional knockout of their proposed upstream TFs. However, even the modest 38-52% reductions of *Lgmn* and *Tlr2* mRNA in the ATF4, ATF3 and/or CHOP gRNA conditions turn out to reflect only a single exceptionally high outlier amongst the non-targeting control samples. That outlier sample, “non.3d.10,” exhibited normalized expression values for *Lgmn* and *Tlr2* that were 2.1-4.9-fold greater than the corresponding values for the other four non-targeting control samples, sufficient to account for the significant effects calculated for the ATF3 gRNA condition. Mild suppression of ATF4/C/EBPγ program gene *Prdx6* by in ATF3, CHOP or CEBPG gRNA conditions is paralleled, incongruously, by a 76% increase in *Prdx6* mRNA upon ATF4 gRNA expression. Moreover, none of these purported “validation” and exemplar genes is significantly different from the uninjured controls at 3dpc in Tian *et al*.’s time course data set (GSE184547), making them unlikely reporters of TF function in the injury response. Together, these observations do not provide evidence to support on-target knockout of their proposed upstream TFs.

**Figure 5.**
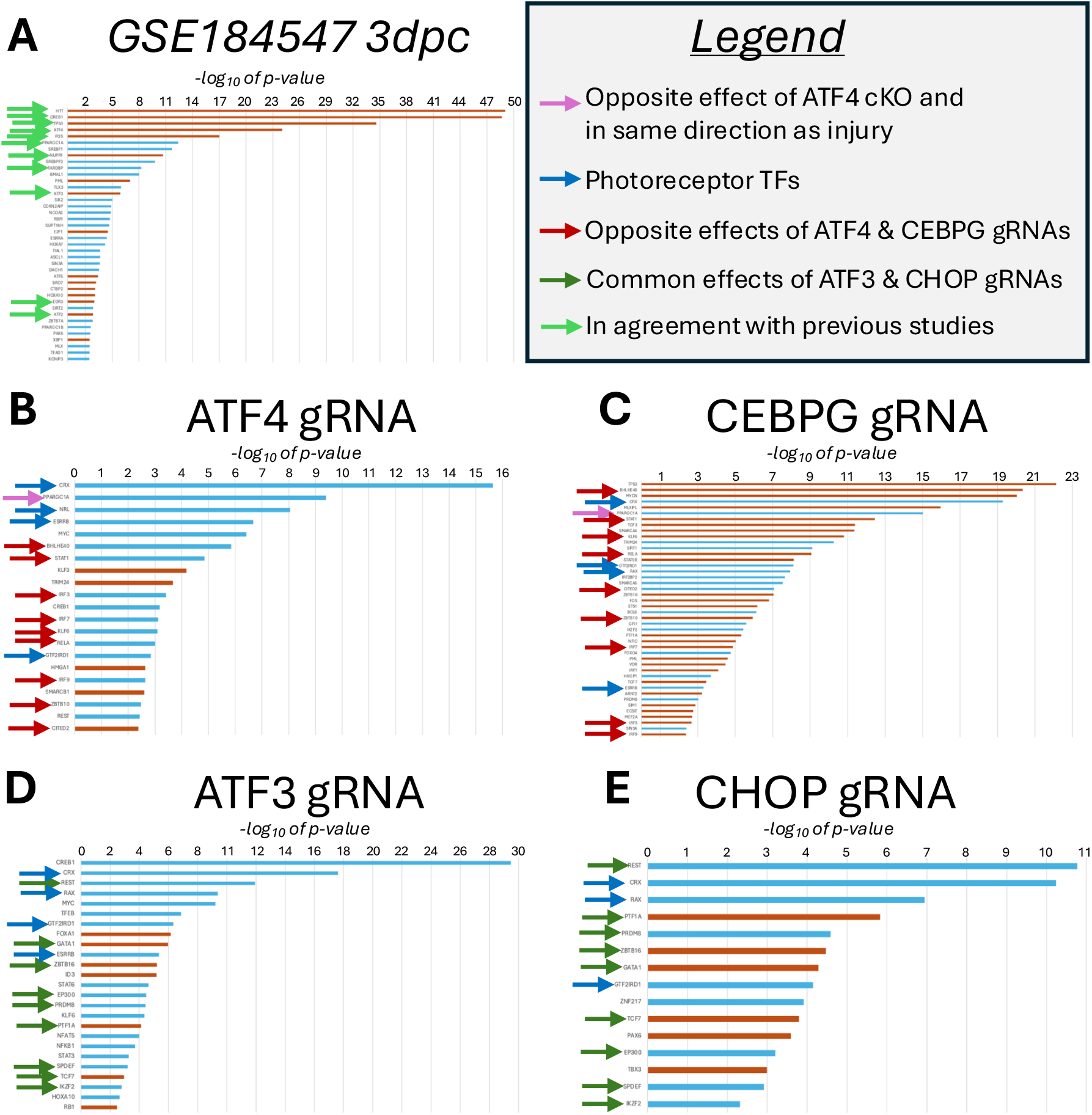
Ingenuity Pathway Analysis suggests a prominent rod photoreceptor gene contribution to differentially expressed genes attributed to gRNA effects. (A) Independent “Upstream Transcription Regulator” analysis of Tian *et al*.’s GSE184547 injury time course data set at 3 days post-crush (3dpc) proposes well-described injury response pathways common to other studies of optic nerve crush, including a prominent role for ATF4 based on upregulation of many of its known target genes. (B-C) “Upstream Transcription Regulator” analysis of Tian *et al*.’s GSE190667 gRNA effects data set at 3dpc identifies entirely distinct sets of potential TF regulators from those inferred for the injury response, with no inference of ATF4 downregulation in response to ATF4 gRNA and anti-correlation between ATF4 and CEBPG gRNAs for multiple inferred regulators. (D-E) Prominent roles for TFs important for photoreceptor development and function, such as CRX and RAX, but not known to be expressed by RGCs, are inferred for ATF3 and CHOP gRNA, similar to what is found with ATF4 and CEBPG gRNA. Burnt orange bars indicate a positive Activation Score inferred by the “Upstream Transcription Regulator” algorithm, and turquoise bars indicate a negative Activation Score.

Ineffective CRISPR-mediated knockout of the targeted TFs offers a likely explanation for many of Tian *et al*.’s perplexing findings but raises another question. In the absence of TF knockout, what is the source of the thousands of reported DEGs, far beyond the number that might be expected from gRNA off-target effects?

### Many differentially expressed genes attributed to TF function instead result from differential RGC enrichment

To begin to address that question, Ingenuity Pathway Analysis (IPA) “Upstream Transcription Regulator” algorithms were applied to two of the published data sets: GSE184547, which reported the injury time course, and GSE190667, which reported the gRNA effects at 3dpc. Analysis of the significant expression changes after optic nerve injury at 3dpc (GSE184547) found notable consistency with other studies of optic nerve crush, with the top six TF regulators proposed by IPA agreeing with a similar analysis in an independent study^8^ (**Figure 5A**). These include inferred activation of ATF3 and ATF4, consistent with the upregulation of well-described ATF4 target genes, including *Chac1, Slc7a5, Phgdh, Sesn2*, among numerous others^8^. Tian *et al*.’s GSE184547 time course data therefore agree with other studies in readily detecting extensive regulation of *bona fide* ATF4 target genes after injury.

In contrast, similar IPA evaluation of the gRNA effects at 3dpc in the GSE190667 data sets reveals little coherence, even within those data. Exceptionally few of the same IPA-determined Transcription Regulators are inversely modulated by gRNA expression, including no suggestion that gene sets regulated by ATF3 or ATF4 were substantially reduced in any gRNA condition (**Figure 5B-E**). Moreover, multiple inferred regulators that are not seen in the injury response, such as the known lymphocyte factors BHLHE40, STAT1, and IRF7^21–23^, exhibit opposite Activation Scores for ATF4 and CEBPG gRNA conditions, with the strange implication that prominent pathways are anti-correlated for these likely heterodimeric TF partners (**Figure 5B-C**). Another conspicuous observation is that negative Activation Scores for CRX and other photoreceptor-enriched TFs, such as RAX, ESRRB, NRL, and GTF2IRD1, are proposed by the IPA algorithm across all four gRNA conditions (**Figure 5B-E**)^24–26^. That result raised the possibility that differential photoreceptor contamination in FACS-enriched RGC samples might have generated numerous DEGs even in the absence of gRNA efficacy. That possibility was made more likely by a reported experimental design (Tian *et al*.’s Table S5A) in which the control non-targeting RGC samples from male mice were generated in distinct batches (“batch 2” and “batch 4”) from the two ATF4 gRNA samples (“batch 1”) and the batch composed of the other three gRNA conditions (“batch 3”, female mice)^2^.

The hypothesis that systematic differences in sorting precision and accuracy can account for the reported DEGs predicts that collections of non-RGC cell type-specific transcripts would exhibit expression patterns that cluster together by sample and by batch. Heat maps were therefore generated of selected transcripts that could serve as cell type markers for rods and other contaminating cell types implied by the Ingenuity Pathway Analysis. As expected, transcripts clustered into groups consistent with their reporting of cell type (**Figure 6A**). Among these was a cluster of genes expressed by macrophages and monocytes, including *Laptm5, Hpgds*, and *Csf1r*, which, like the macrophage marker *Lgmn*, were strongly enriched only in the lone “batch 4” non-targeting control sample^27–31^. Notably, related expression patterns were shared by all eight of the program genes purported to indicate positive regulation of the highlighted “MAPK Signaling / Neuroinflammation” and “TLR Signaling and Processing” pathways by ATF3 in Tian *et al*.’s revised Figure 5A (*Dusp1, Axl, Jun, Ccl2, Cd86, Tlr2, Nfkbia, Lgmn*). An additional cluster of T cell markers, including T cell receptor alpha constant (*Trac*), T cell receptor β constant 2 (*Trbc2*), and *Cd4*, was especially enriched in CEBPG gRNA samples relative to CHOP and ATF3 gRNA samples, suggestive of systematic sorting differences among distinct conditions even within “batch 3.” Consistent with that possibility, non-RGC transcripts accounted for significant DEGs between the ATF3 gRNA and CHOP gRNA conditions (**Supplemental Figure S1A-C**). The sharpest cluster, consistent with the Ingenuity Pathway Analysis, was a collection of rod photoreceptor markers, such as rhodopsin (*Rho*), retinitis pigmentosa 1 (*Rp1*), recoverin (*Rcvrn*) and phosducin (*Pdc*), which were enriched ∼7-12-fold only in the “batch 2” non-targeting control samples (**Figure 6A**). Notably, *Atf4*, a core TF, and *Mef2c*, an exemplar of the “Common” transcription program, exhibit similar patterns of expression, suggesting that the apparent reduction of these transcripts across all four gRNA conditions largely reflects their enrichment in rod photoreceptors rather than effects of knockout^32^. Contrasting with these batch and condition effects in the GSE190667 RNA-seq data set, variance in rod photoreceptor and immune cell markers in Tian *et al*.’s separate GSE184547 injury time course data set is not generally systematic across time points (**Figure 6B**). For example, high rod contamination in one control sample (“no.crush.rep1”) was offset by low rod presence in another control sample (“no.crush.rep3”), consistent with the absence of rod-specific TFs amongst the Upstream Regulators proposed by IPA (**Figure 5A**). Additional heat maps reveal that differential cell sorting contributed substantially to the genes included in all three of the revised Table S6 transcriptional programs (**Figure 6C-E**).

**Figure 6.**
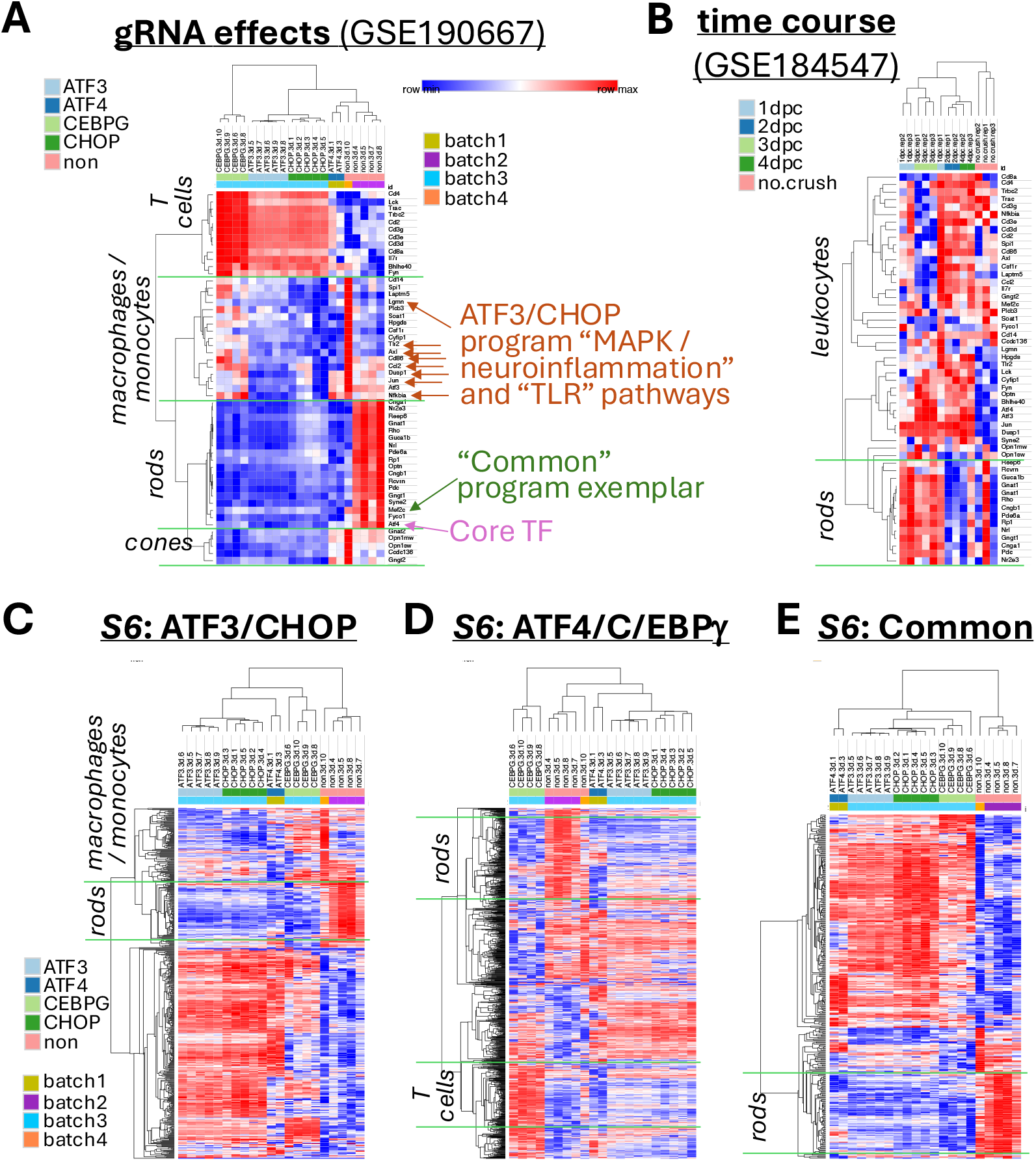
Proposed “core transcription programs” include genes from differential RGC enrichment across conditions. (A-B) Heat map of expression of cell type marker genes for rod and cone photoreceptors and various immune cells, along with genes presented in Tian *et al*.’s Figure 5A to support prominent roles for the ATF3/CHOP program in pathways of innate immunity (*brown arrows*), the core TF gene *Atf4* (*pink arrow*), and the “Common” program exemplar gene *Mef2c* (*dark green arrow*). (C-E) Clusters of genes consistent with rods, T cells, and macrophages/monocytes are found within each of the three proposed programs found in Tian *et al*.’s Table S6. All data are from independent DESeq2 analyses of Tian *et al*.’s GSE184547 (injury time course) and GSE190667 (gRNA effects) RNA-seq data sets.

**Figure 7.**
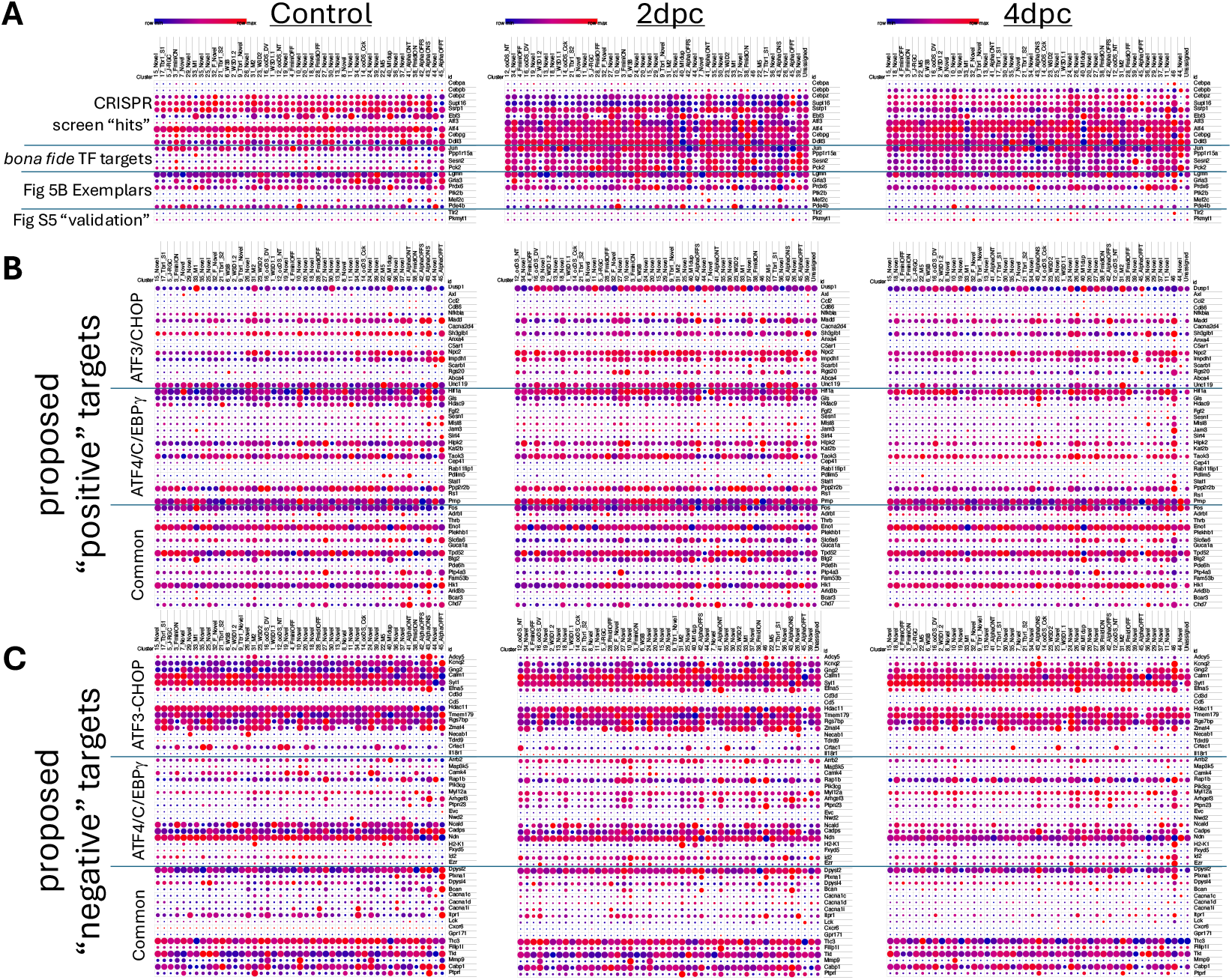
Publicly available scRNA-seq data reports that numerous proposed program genes are not expressed in RGCs or are not injury-responsive. (A) scRNA-seq data from injured (2 and 4 days post-crush) and uninjured (Control) RGC subtypes for prominent genes relevant to the Tian *et al*. manuscript’s conclusions: ten reported “hits” from that study’s comprehensive *in vivo* CRISPR screen (including the four “core” TFs); known targets of ATF3 and ATF4; genes presented as exemplars of the proposed “core transcription programs” in Tian *et al*.’s Figure 5B; and proposed program genes purported to be validated by IHC in Tian *et al*.’s Figure S5. (B) scRNA-seq for proposed “positive” targets across all three putative programs, including genes from Tian *et al*.’s Figure 5A and top DEGs from Tian *et al*.’s Table S6. (C) scRNA-seq for proposed “negative” targets across all three putative programs, including genes from Tian *et al*.’s Figure 5A and top DEGs from Tian *et al*.’s Table S6. All data are from Tran *et al*. (2019).

These analyses suggest that many of the proposed program genes are not expressed by uninjured or injured RGCs. To further examine that possibility, an independent scRNA-seq resource was consulted that reported the RGC subtype-specific expression changes over time following optic nerve crush, including at 2dpc and 4dpc^11^. Plots were generated from this publicly available resource for multiple classes of genes. The first of these plots consisted of: (1) genes reported as CRISPR screening “hits” in Tian *et al*.’s Figure 1; (2) *bona fide* TF target genes; (3) the six exemplar program genes of Tian *et al*.’s Figure 5B; and (4) the two IHC “validated” genes of Tian et al.’s Figure S5 (**Figure 6A**). Whereas *bona fide* stress-responsive TF targets *Jun, Ppp1r15a, Sesn2*, and *Pck2* exhibited the expected injury-induced upregulation, several other transcripts were not reported by scRNA-seq data set to be expressed by RGCs. These include the reported CRISPR screen “hit” *Cebpa*, the ATF4/CEBPG program exemplar *Ptk2b*, the “Common” program exemplar *Mef2c*, and the two IHC “validation” genes *Pkmyt1* and *Tlr2*. This latter finding is particularly significant. The PKMYT1 IHC referred to in the text of the Tian *et al*. article is not found in the published Supplemental data. The IHC for TLR2 in Tian *et al*.’s Figure S5 is therefore the only reported validation of a proposed program gene in that manuscript. Previous studies of TLR2 in retina following optic nerve crush reported its expression in astrocytes, Müller glia, and macrophages^33^. In the IHC of Tian *et al*., however, TLR2 protein appeared to exhibit ∼10-fold upregulation in the ganglion cell layer at 3dpc that largely overlaps with the RGC marker RBPMS. The purported TLR2 IHC signal is shown to have been completely and consistently suppressed by ATF3+CHOP gRNA but not by ATF4+CEBPG gRNA. Those observations are incompatible with the apparent lack of *Tlr2* mRNA expression by RGCs^11^. Moreover, the bulk RNA-seq analyses of Tian *et al*. indicate that the apparent reduction of *Tlr2* mRNA upon gRNA expression reflects instead only the exceptionally high contamination by TLR2-expressing immune cells in the lone “batch 4” non-targeting control sample. The provided IHC data is therefore unlikely to reflect underlying TLR2 protein in RGCs and do not serve as effective validation of the proposed ATF3/CHOP-dependent program.

Additional scRNA-seq plots reveal that many other program genes reported in Tian *et al*.’s Figure 5A and Table S6 also are not substantially expressed by RGCs or are not modulated by injury within RGCs (**Figure 6B-C**). Notably, these include genes that underlie Tian *et al*.’s central claim that the ATF3/CHOP program regulates genes related to innate immunity, such as the AXL receptor tyrosine kinase (*Axl*), the co-stimulatory molecule B7.2 (*Cd86*), and the monocyte chemoattractant protein-1 (MCP-1 / *Ccl2*), that, along with *Tlr2*, are highly expressed by antigen-presenting cells^34–37^. Similarly, minimal RGC expression of genes like *Fgf2, Sesn1* and *Mlst8* undermine support for the proposed activation of “intrinsic neuronal stressors” like “Autophagy” by the ATF4/C/EBPγ program.

This evaluation of the published findings suggests that the majority of DEGs attributed to TF function instead reflect differential enrichment of RGCs across batches without adequate distribution of conditions across batches to mitigate this confound. These results provide clarity for how ineffective gRNA knockout could nevertheless generate DEGs to populate the proposed neurodegenerative transcription programs. The reported findings using these gRNAs, however, do not substantially inform the functions of the targeted TFs in neurodegeneration.

## Discussion

Deciphering the transcription factors (TFs) and programs that underlie neurodegeneration and axon regenerative potential following CNS axon injury may generate opportunities for neuroprotection and CNS repair. As the most comprehensive investigations of these questions, the study reported by Tian *et al*. (2022) offered to define the “core” TFs for neurodegeneration, their programs, and their relationships, providing the foundation for numerous potential research and therapeutic directions. A second Correction refined these proposed programs based in part on greater statistical stringency for one of the criteria used to define the programs. Surprisingly, however, the analysis reported here finds that this increased stringency was accompanied not just by a contraction of genes previously included in the programs but also addition of numerous genes that failed to meet essential criteria of the originally proposed programs. This *Matters Arising* evaluation of the Correction reveals that these fundamentally redefined “programs” demonstrate little coordination between pair of core TFs, rendering a central premise of the original manuscript to be insubstantial. Only four genes of the ATF3/CHOP program are depicted in Tian *et al*.’s revised Figure 5A as co-regulated by ATF3 and CHOP, two of which (*Axl, Cd86*) appear not to be expressed by RGCs. Similarly, only four genes of the revised ATF4/C/EBPγ program are depicted as co-regulated by ATF4 and C/EBPγ, and one of those (*Prdx6*) is reported to exhibit opposite effects of ATF4 and CEBPG gRNA. This latter result is particularly surprising, as even moderate knockout of these TFs could be expected to uncover some of the numerous injury-induced genes known to be bound and regulated by the ATF4-C/EBPγ heterodimer. Such findings raise doubt as to whether the most essential tool of the study, CRISPR-mediated knockout, was sufficiently effective to be of value in informing the roles of these TFs in injury-induced neurodegeneration.

It is difficult to reconcile the apparent absence of functional knockout of ATF4, CEBPG, and CHOP demonstrated here (and the seeming moderate knockout of ATF3) with the exceptionally robust reduction in their protein levels implied by immunohistochemistry (IHC) in Tian *et al*.’s Figures 6B-E and S6D-G. The reported IHC results for TLR2, which previous studies have found not to be expressed by injured or uninjured RGCs, raise the possibility that findings of this pattern may be common with the immunostaining techniques used. IHC for ATF3, robust across labs, is typically tightly nuclear^38–40^, in possible contrast to Tian *et al*.’s Figure 6B, whereas it can be challenging to validate endogenous ATF4, CHOP, and C/EBPγ immunostaining in this and other systems. In the absence of additional validation, the minimally variant ∼8-15-fold reported reductions in these proteins in Tian *et al*.’s Figures 6B-E and S6D-G may best be interpreted with caution, as those reductions appear to far exceed the knockout eficiencies of ∼65-75% reported in Tian *et al*.’s Figure S1B-E for the proof-of-principle targets SATB1 and PTEN.

The current *Matters Arising* analysis highlights the importance of batch design in transcriptomic studies, especially those that rely on bulk RNA-seq of enriched cell populations. Though FACS and other cell type enrichment techniques are necessarily imperfect, the consequences of such imperfections can be substantially mitigated when they are distributed similarly across conditions, as demonstrated by the similarities in the reported post-injury time course (Tian *et al*.’s Table S2 and GSE184547) to the findings of many related studies. However, systematic differences across batches and conditions in non-RGC contamination seem to account for most of the reported gene expression changes and pathways attributed to gRNA-mediated TF knockout (Tian *et al*.’s Table S5 and GSE190667). The current analysis therefore clarifies two unexpected conclusions put forth in the abstract of Tian *et al*.: (1) that the “intrinsic neuronal stressors” of the ATF4/C/EBPγ program are quite distinct from the injury-activated pathways well-described to be driven by these TFs, and (2) that an ATF3/CHOP program regulates “pathways activated by cytokines and innate immunity.” For both primary conclusions, many of the transcripts reported as support for these claims are not expressed by RGCs and/or track with markers of other cell types known to express those genes at high levels.

The above analyses demonstrate that lack of TF knockout, combined with systematic contamination by non-RGCs across conditions, are likely to have generated the strikingly novel findings for the “core transcription programs” underlying injury-induced RGC neurodegeneration reported in Tian *et al*. These results argue against drawing firm conclusions from the published data regarding the seemingly similar, non-additive neuroprotective effects of ATF3 and CHOP gRNAs or the seemingly similar, non-additive neuroprotective effects of ATF4 and CEBPG gRNAs. Though conditional knockout of ATF4 has independently been demonstrated to provide partial RGC neuroprotection^7,8^, it is not difficult to confidently attribute the comparable effects of ATF4 gRNA reported by Tian *et al*. to an equivalent reduction in ATF4 function, suggesting alternative mechanisms for the reported neuroprotection. And though it is plausible that C/EBPγ could play an overlapping role with ATF4 in neurodegeneration, that question has not been adequately resolved by the Tian *et al*. study or its Corrections. The revised data provided by Tian *et al*. therefore are not sufficient to support the proposed “core transcription programs” of ATF3/CHOP and ATF4/C/EBPγ and do not appreciably inform the relationships and functions of these TFs in neurodegeneration.

## Methods

### RNA-seq analysis

Independent DESeq2 analyses of GSE184547 (optic nerve crush injury time course), GSE190667 (effects of gRNA treatments at 3 dpc) was performed using the publicly available OneStopRNA-seq resource by uploading FASTQ files directly from GEO^41^. To maintain consistency with Tian *et al*.’s analyses of differential expression in GSE184547, 2dpc.rep3 and 4dpc.rep1 were excluded. Heat maps of DESeq2 normalized expression values, adjusted to log_2_ and hierarchically clustered by the one minus Pearson correlation metric, were generated using a publicly available resource: Morpheus, https://software.broadinstitute.org/morpheus

### Quantification of overlap between original and revised “core transcription programs”

The original Tian *et al*. Figure 5 and Table S6 were downloaded from the *Neuron* website prior to the replacement of those files with revised versions upon the more recent of the published Corrections. These and other pre-Correction files remain available for download in the PubMed Central version of the manuscript (https://pmc.ncbi.nlm.nih.gov/articles/PMC9391318/). Each row of the original and revised Tables S6 depicts one DNA footprinting-inferred binding site of the proposed “core transcription programs,” with some sites and genes represented on multiple rows, one for each TF-binding site interaction. The number of unique genes represented in each program were counted, along with the number of program TFs by which each gene was inferred to be bound. “Overlapping” DNA footprinting-inferred binding sites were defined as sites listed in Table S6 targeted by two of more “core TFs” within 4 bases of one another, with the large majority within 0-2 bases. Venn Diagrams were generated using the publicly available DeepVenn resource^42^.

### Correlation analyses

Using differential expression values from GSE190667_rna.DEX and independent DESeq2 analyses of GSE190667 and GSE184547 data sets, the VLOOKUP function of Microsoft Excel was used to align reported values for each transcript across data sets, generating new tables and graphs in which the differential expression values (log_2_ fold-change, or logFC) and significance values (false-discovery rate, FDR, or adjusted *p*-value) for each transcript from one data set were plotted against the corresponding values from the related data set. Transcript names were color-coded and highlighted based on their representation in Figure 5A of the more recent Correction^1^, which, for SIRT4 and EFNA5, differs from their representation in the replacement Figure 5 in the parent manuscript^2^.

### Ingenuity Pathway Analysis

Upstream Transcription Regulator analysis of Ingenuity Pathway Analysis (IPA) was applied to top DEGs from the revised GSE190667_rna.DEX and GSE184547_rna.DEX files posted to GEO by authors of Tian *et al*.^1,2,43,44^ after filtering out low expressors (average log_2_ normalized expression<2.5). For ATF3, ATF4, and CHOP gRNA conditions, |logFC|>0.5 and FDR<0.1 cutoffs generated 1635, 1479, and 1535 transcripts for analysis, respectively. For the CEBPG gRNA condition, |logFC|>0.65 and FDR<0.1 cutoffs generated 2934 transcripts for analysis.

### scRNA-seq plots

Publicly available scRNA-seq data from Tran *et al*. (2019) were accessed and plotted using the Single Cell Portal^11,45^: https://singlecell.broadinstitute.org/single_cell/study/SCP509/mouse-retinal-ganglion-cell-adult-atlas-and-optic-nerve-crush-time-series/.

## Data availability

As a *Matters Arising* analysis of independently published work, primary data used in this analysis and the manuscripts that it addresses may be available upon request to the corresponding author of Tian *et al*. (2022). In addition to accessing the GSE190667 and GSE184547 data sets through OneStopRNA-seq as described above, the present work relied on the following resources publicly available at the time of submission: the revised spreadsheet GSE190667_rna.DEX; Tian *et al*.’s revised Figure 5, revised Table S5, revised Table S6, and Table S3. Other resources utilized in this assessment include Tian *et al*.’s original Figure 5, original Table S6, and original Table S2, which were, atypically, removed from the published manuscript upon posting of the Corrections, though they remain available in the PubMed Central version of the manuscript.

## Figure legends

**Supplemental Figure S1.**
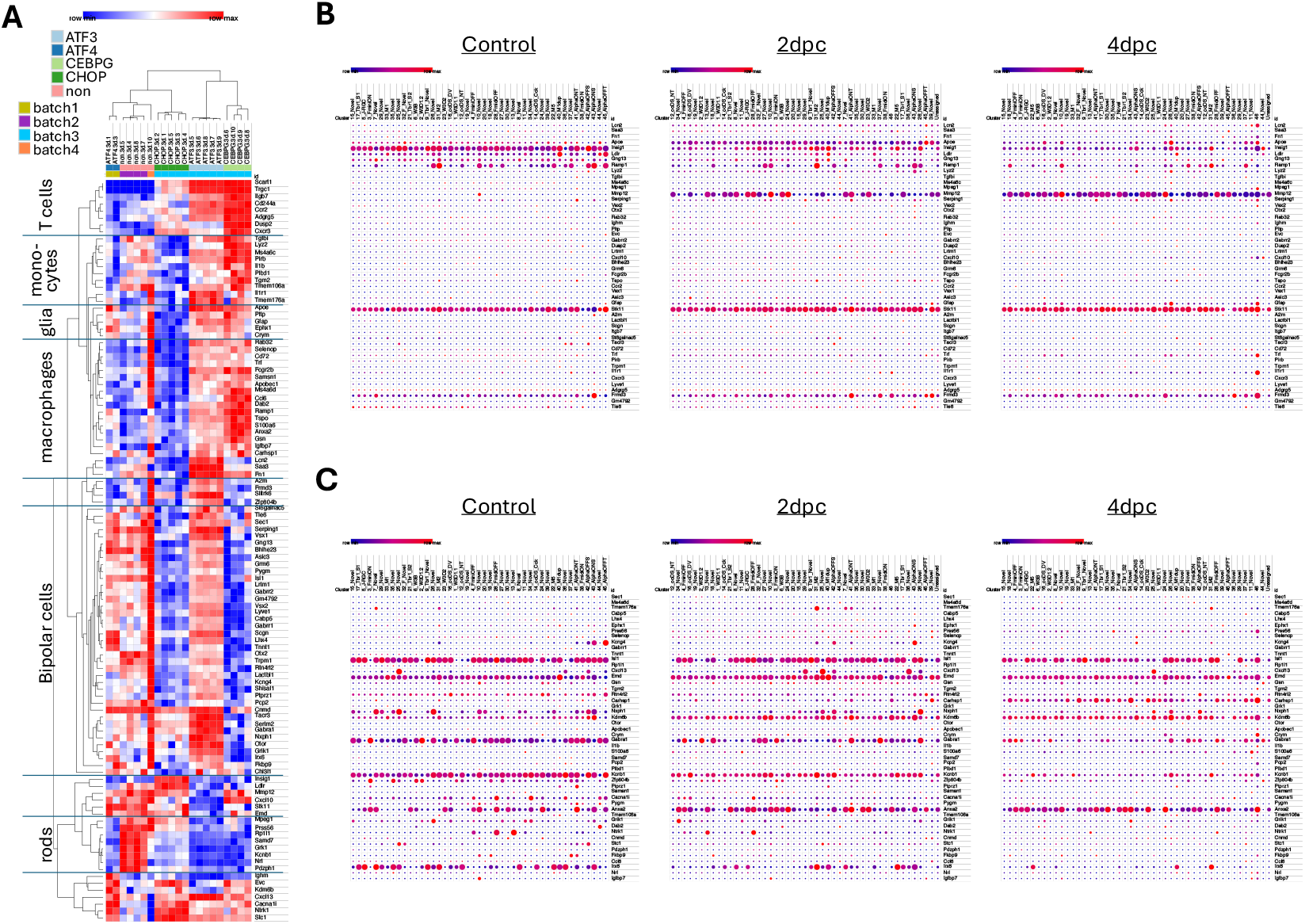
Differential cell sorting account for differences between ATF3 gRNA and CHOP gRNA effects. (A) Heat map of differentially expressed genes between ATF3 gRNA and CHOP gRNA conditions (*adj. p*<0.1, |logFC|>1, basemean>500) after hierarchical clustering, annotated with cell types known to express similar clusters of genes. Data are from an independent DESeq2 analysis of Tian et al.’s GSE190667 (gRNA effects) RNA-seq data set. (B-C) Publicly available scRNA-seq data reports few of these genes to be also expressed by injured (2dpc, 4 dpc) or uninjured (Control) RGCs. Data are from Tran *et al*. (2019).

